# Epithelial Infection with *Candida albicans* Elicits a Multi-system Response in Planarians

**DOI:** 10.1101/2020.11.12.380519

**Authors:** Eli Isael Maciel, Ashley Valle Arevalo, Benjamin Ziman, Clarissa J. Nobile, Néstor J. Oviedo

**Author notes:** To whom correspondence should be addressed and, Department of Molecular & Cell Biology, University of California, Merced. 5200 North Lake Road, Merced, CA 95343. These authors contributed equally to this work (Eli Isael Maciel, Ashley Valle Arevalo). **Summary Statement:** Fungal infection elicits a multi-system innate immune response in planarians involving stem cells, the excretory system, and the nervous system.

## Abstract

*Candida albicans* is one of the most common fungal pathogens of humans. Prior work introduced the planarian *Schmidtea mediterranea* as a new model system to study the host response to fungal infection at the organismal level. In the current study, we analyzed host-pathogen changes that occurred *in situ* during early infection with *C. albicans*. We found that the transcription factor Bcr1 and its downstream adhesin Als3 are required for *C. albicans* to adhere to and colonize the planarian epithelial surface, and that adherence of *C. albicans* triggers a multi-system host response that is mediated by the Dectin signaling pathway. This infection response is characterized by two peaks of stem cell divisions and transcriptional changes in differentiated tissues including the nervous and the excretory systems. This response bears some resemblance to a wound-like response to physical injury; however, it takes place without visible tissue damage and it engages a distinct set of progenitor cells. Overall, we identified two *C. albicans* proteins that mediate epithelial infection of planarians and a comprehensive host response facilitated by diverse tissues to effectively clear the infection.

## INTRODUCTION

The innate immune system is the first line of defense against invading pathogens (Espinosa and Rivera, 2016; Peiris et al., 2014). The complex interactions between the pathogen and the host is a constantly evolving arms race, where the pathogen aims to invade and proliferate in the host, while the host aims to defend against infection. Both the host and pathogen responses to one another are known to be influenced by the physical presence of the infecting microorganism on specific host tissues, by the production of pathogen-associated virulence factors, and by the extent of host tissue damaged during the infection. Nonetheless, the underlying molecular mechanisms associated with these host-pathogen responses are largely unknown.

*Candida albicans* is an opportunistic human fungal pathogen that can cause both superficial mucosal and life-threatening systemic infections, particularly in immunocompromised individuals (Filler, 2013; Pellon et al., 2020). *C. albicans* efficiently adapts to the dynamic host environment by quickly responding to host environmental cues, such as contact with epithelial cells, temperature, pH, nutrients, and iron levels (Brunke and Hube, 2014). *C. albicans* possesses a number of virulence traits that allow this pathogen to evade host innate immune defenses, such as the ability to change morphologies from the round budding yeast-form to the elongated hyphal form (Braun and Johnson, 1997; Mitchell, 1998; Saville et al., 2003). The yeast to hyphal transition is critical for *C. albicans* to invade and damage host tissues and to disseminate into the bloodstream (Wilson et al., 2016). Additionally, *C. albicans* can adhere to and colonize host tissues by forming recalcitrant biofilms on mucosal surfaces that provide physical protection from the host innate immune response (Nobile and Johnson, 2015; Valle Arevalo and Nobile, 2020).

*C. albicans* uses two main strategies to invade and damage host epithelial cells. The first involves inducing endocytosis by host cells, while the second involves active physical penetration of host cells by *C. albicans* hyphal cells (Filler, 2013; Filler et al., 1995; Pellon et al., 2020; Wachtler et al., 2012). Adhering to epithelial cells and forming hyphae is the first step for *C. albicans* to invade host cells via either induced endocytosis or active physical penetration strategies (Wilson et al., 2016). For induced endocytosis, the process is mediated by fungal cell surface proteins (e.g. Ssa1 and Als3) that bind to cadherins on host cell surfaces (Filler, 2013; Moreno-Ruiz et al., 2009; Nobile et al., 2006; Phan et al., 2007; Sun et al., 2010). Active physical penetration of host cells by *C. albicans* cells, on the other hand, is a fungal-driven process that relies on the ability of hyphal cells to penetrate the epithelial cells without host activity (Wilson et al., 2016). Overall, the molecular mechanisms mediating both strategies are not fully understood.

To date, the analysis of host-pathogen interactions, including our understanding of pathogen invasion strategies, has been possible through the use of *in vitro*, *ex vivo*, and *in vivo* infection models. Work in *C. albicans* has led to the identification of several molecular players and signaling pathways involved in the host-pathogen response to acute *C. albicans* infections (Pellon et al., 2020). Nevertheless, the host response to infection likely goes beyond these localized interactions with the pathogen and may involve complex long-range intercellular communication between host tissues and organs to orchestrate an effective response against the invading pathogen (Godinho-Silva et al., 2019). Visualizing the complexity of the host response in real-time and integrating the contribution of different organs and systems has been challenging with currently available animal models (Bergeron et al., 2017; Chamilos et al., 2007; Glittenberg et al., 2011; Gratacap et al., 2017; Mallick et al., 2016; Mylonakis, 2008; Mylonakis et al., 2007; Peterson and Pukkila-Worley, 2018; Pukkila-Worley et al., 2009; Segal and Frenkel, 2018).

Previously, we introduced the planarian *Schmidtea mediterranea* as an *in vivo* infection model to study host-pathogen interactions with *C. albicans* (Maciel et al., 2019). We found that planarians can be conveniently infected by adding *C. albicans* cells directly into the media where planarians live, meaning that infection does not require invasive procedures, such as injections. This “passive” infection method allows for the spatiotemporal tracing of *C. albicans* throughout the course of an infection, beginning with its initial interaction with the planarian epithelial surface. The aggressive invasiveness of *C. albicans* hyphal cells on planarian tissues is counteracted by the planarian innate immune response that eventually completely clears the infection in about a week (Maciel et al., 2019). Aside from prompting an innate immune response, infection of planarians with *C. albicans* triggers a mitotic response that is mediated by pluripotent adult stem cells known as neoblasts, the only somatic cell population in planarians with the capacity to divide (Maciel et al., 2019). This finding is intriguing because fungal infections have also been shown to activate mammalian stem cell proliferation, which is believed to potentiate the host innate immune response to infection (Megías et al., 2016; Megías et al., 2012; Yáñez et al., 2011; Yáñez et al., 2009; Yang et al., 2012). These findings suggest the possibility of evolutionary conservation of the host stem cell and innate immune responses between mammals and planarians. Nonetheless, the underlying mechanisms used by planarians to effectively clear *C. albicans* infections remain unclear.

In the present study, we show that infection of planarians with *C. albicans* triggers systemic waves of neoblast proliferation that can be detected within six hours post-infection. These increases in neoblast proliferation are preceded by the adherence of *C. albicans* cells to the host epithelial cell surface. Furthermore, the presence of *C. albicans* elicits a reaction in planarians that resembles a regenerative response, including an increase in expression of the early growth response gene *Smed-egr2 (egr2)*, and an immediate wound response via the ERK signaling pathway (Owlarn et al., 2017; Wenemoser et al., 2012; Wurtzel et al., 2015). We also found that the host response is mediated by the *C. albicans* transcriptional regulator Bcr1 and its downstream cell surface protein Als3, which are key players in fungal cell adherence.

Finally, we also observed that upon infection of planarians with *C. albicans*, there is an increase in expression of components of the Dectin signaling pathway within the host innate immune system, and an increase in the expression of host markers of neoblast subpopulations as well as markers of the nervous and excretory systems. Together, our findings demonstrate that the initial adherence of fungal cells to the epithelial surface of the planarian host elicits a multi-system response in the host involving the innate, excretory, and nervous systems, as well as neoblast function.

## RESULTS

### Adherence of C. albicans to the planarian epithelial surface initiates an early wound-like response

To assess the host response to *C. albicans* infection, planarians were infected with a lethal concentration (25 ×10^6^ cells/mL) of *C. albicans* cells via soaking as previously described (Maciel et al., 2019). Our prior results demonstrated that planarians responded to fungal infection with an increase in mitotic activity; however, it was unclear when during the infection this fungal-induced proliferative response was occurring (Maciel et al., 2019). To address this, we infected planarians with *C. albicans* and analyzed host mitotic activity temporally over the course of 48 hours (**Figure 1A-B**; see **Supplemental Figure 1** for descriptions of the spatial orientations of the planarian whole mounts). We observed no host behavioral or macroscopic differences within the first 48 hours post-infection (hpi) between the uninfected and experimental groups. We did, however, detect two host mitotic peaks occurring at six and 24 hpi, which were preceded by decreased mitotic activity (**Figure 1B**). The largest burst in mitotic activity (~150%) was observed six hpi, and was spatially increased throughout the planarian body, suggesting a system-wide neoblast response (**Figure 1A-B**).

**Figure 1:**
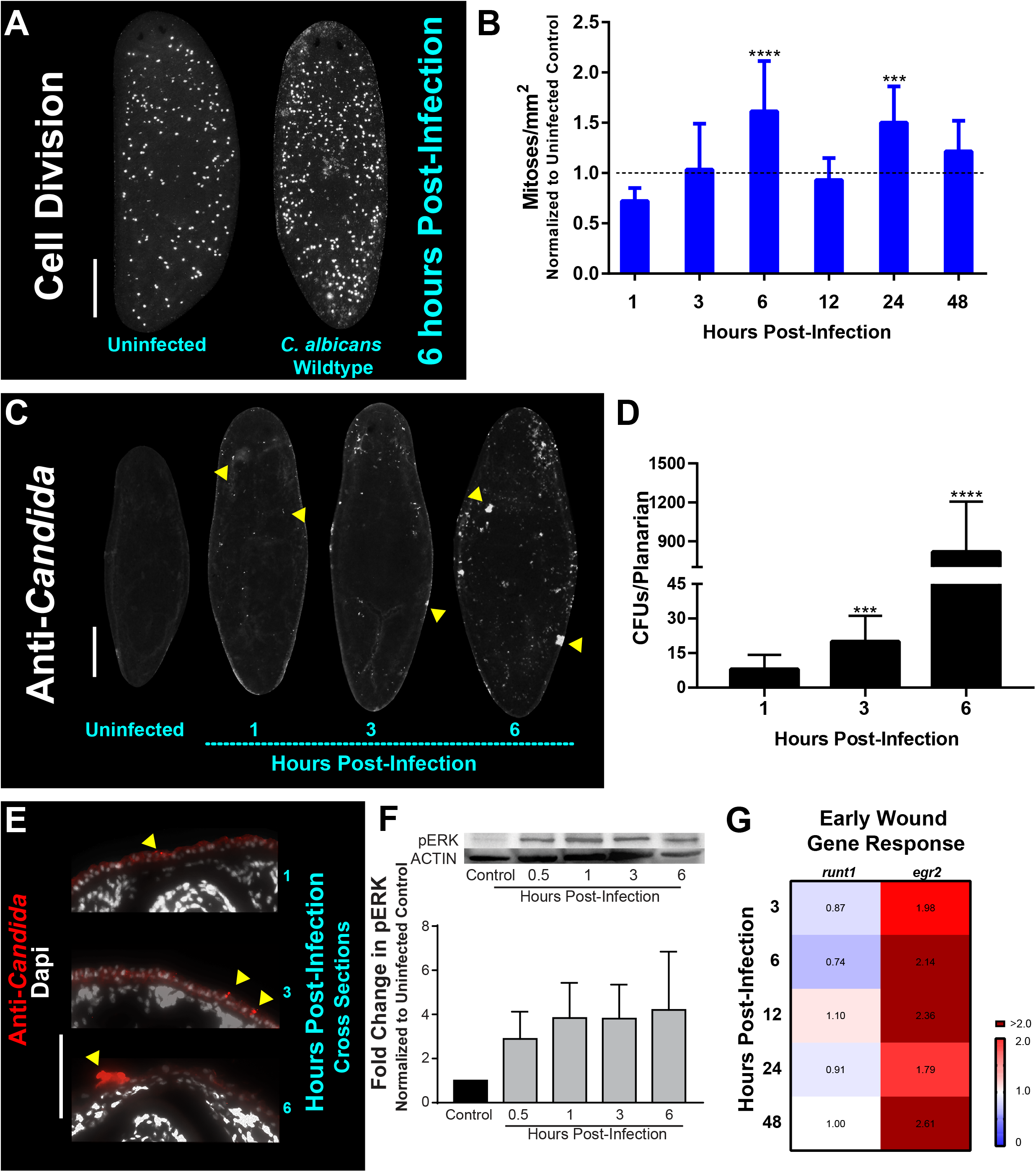
*C. albicans* infection leads to an early wound and mitotic response in planarians. A) Whole-mount immunostaining with anti-phospho-histone H3 (ser10) antibody for planarians, which labels mitotic cells (white foci) at six hours post-infection with the *C. albicans* wildtype strain. B) Number of mitoses in response to *C. albicans* cells between one to 48 hours normalized to an uninfected planarian. C) Whole-mount anti-*Candida* antibody stain at one, three, and six hours post-infection (white foci). D) *C. albicans* colony forming units (CFUs) normalized per planarian. Ten animals per infection timepoint were used. E) Transverse cross-section images from the dorsal side of the planarian body showing the adherence of the *C. albicans* wildtype strain (red signal) at one, three, and six hours post-infection. F) Western blot of planarian phosphorylated-ERK (p-ERK) and quantification at different timepoints of a *C. albicans* infection. β-tubulin was used as an internal control. G) Gene expression levels represented in a heat map of wound-induced response planarian genes *runt1* and *egr2* at the different infection timepoints. Gene expression is represented by fold change normalized to an uninfected control. Color scale depicts red as upregulation and blue as downregulation; burgundy demonstrates upregulation greater than twofold. Data were obtained in triplicate per experiment for at least two biological and technical replicates. *C. albicans* infections were performed using 25 million cells/mL. All graphs represent mean ±SEM. Statistical comparisons are against the uninfected control. Scale bar is 200μm. Two-way ANOVA, *P<0.01; **P<0.005; ****P<0.0001.

To determine if the mitotic response was due to the adherence or penetration of *C. albicans* cells through the epithelial surface of planarians, we focused on early timepoints of one, three, and six hpi. These experiments revealed that as early as one hpi, *C. albicans* cells began to adhere to the planarian epithelial surface and that by six hpi, the fungal cells were readily clustered throughout the epithelial surface (**Figure 1C**). The increased presence of *C. albicans* cells on the planarians during these early timepoints was determined by macerating planarians and measuring colony forming units (CFUs) after plating the homogenized slurry on agar plates. We found that in the first six hpi, the number of *C. albicans* cells increased by two orders of magnitude (i.e. ~10 CFUs at one hpi compared to ~1,000 CFUs at six hpi) (**Figure 1D**). To discern if the increase in *C. albicans* cells during this early infection was limited to the superficial adherence of *C. albicans* cells to the host epithelial surface or to penetration of *C. albicans* cells into host tissues, we performed immunostaining on transverse cross sections using an antibody specific for *C. albicans*. We found that *C. albicans* cells were adhered to the epithelial surface of the planarians, while no *C. albicans* antibody signal was detected in deep tissues even after six hpi (**Figure 1E**). This finding is consistent with previous results demonstrating that *C. albicans* cells attach to the host epithelial surface during early timepoints of infection (Maciel et al., 2019).

In response to physical injury, planarians have been shown to elicit a regenerative mitotic response occurring six hours post-injury (Saló and Baguñà, 1984; Wenemoser and Reddien, 2010). Due to the apparent overlap between injury stimuli (i.e. the presence of penetrating *C. albicans* hyphal cells) and the mitotic response observed during the early infection period (six hpi), we hypothesized that invading *C. albicans* hyphal cells could trigger a wound response in planarians during early infection.

In response to injury in planarians, the planarian extracellular regulated kinase (ERK) becomes activated via phosphorylation (pERK), and is among the earliest events essential for regeneration initiation, occurring ~15 minutes post-injury (Owlarn et al., 2017). We, therefore, measured pERK protein levels at early infection timepoints using a western blot and found that there was a gradual increase in pERK levels, as early as 30 minutes post-infection (~threefold increase), which increased to about fourfold by six hpi (**Figure 1F**). We also measured the post-infection expression levels of *Smed-runt1 (runt1)* and *Smed-egr2* (*egr2*), two early wound response genes in planarians (Owlarn et al., 2017; Sandmann et al., 2011; Wenemoser et al., 2012; Wurtzel et al., 2015). We observed an increase in the expression of *egr2* (~twofold) over the first 48 hpi (**Figure 1G**). Intriguingly, *runt1* was either slightly reduced or showed no changes in expression over the first 48 hpi (**Figure 1G**). Overall, these results suggest that the adherence of *C. albicans* cells to the epithelial surfaces of planarians shortly after infection triggers a regenerative wound-like response in the host.

### Als3 and Bcr1 are required for C. albicans to adhere to and colonize the planarian epithelial surface

Next, we sought to test if the dynamics of infection in planarians can be modulated by genetically disrupting *C. albicans* virulence factors known to contribute to disease progression in the host (Mayer et al., 2013). Based on our observations that *C. albicans* adheres to planarians in order to invade them, we decided to focus our attention on *C. albicans* virulence factors associated with adherence and biofilm formation. The agglutinin-like sequence (Als) family of cell surface glycoproteins are important *C. albicans* adhesins that mediate cell-cell and cell-substrate adherence (Hoyer and Cota, 2016). Two major Als proteins that are known to be important for adherence and biofilm formation are Als1 and Als3 (Nobile et al., 2006; Nobile et al., 2008). In addition to these two adhesins, a known master regulator of *C. albicans* biofilm formation that controls the expression of *ALS1* and *ALS3* is Bcr1 (Nobile et al., 2006; Nobile et al., 2012; Nobile and Mitchell, 2005). We, therefore, assessed the ability of the *als1*, *als3*, and *bcr1* mutant strains to infect planarians. In addition to adherence, *C. albicans* also produces secreted aspartyl proteases (Saps) that are important virulence factors in causing host cell damage (Mayer et al., 2013; Naglik et al., 2003). We, therefore, also assessed the ability of two triple deletion *SAP* mutant strains (encompassing the major *SAPs* involved in virulence), the *sap1/sap2/sap3* and the *sap4/sap5/sap6* mutant strains, to infect planarians.

To our surprise, we found that all *C. albicans* mutant and wildtype strains were capable of adhering to the planarian epithelial surface. Qualitatively, it appeared that the *als3* and *bcr1* mutant strains displayed less adherence six hpi (**Figure 2A**). Consistent with this observation, our results revealed an increase in the survival of planarians when they were infected with lethal doses of the *als3* and *bcr1* mutant strains compared to the wildtype strain (**Figure 2B**). Additionally, we discovered that when the *C. albicans* wildtype strain came into contact with planarians, there was a dramatic increase in the gene expression levels of *ALS1, ALS3*, and *BCR1* (**Figure 2C**). Overall, these results suggest that fungal cell adherence mediated by the adhesin Als3 and the transcription factor Bcr1 is important for *C. albicans* to colonize the epithelial surface of planarians during an infection.

**Figure 2:**
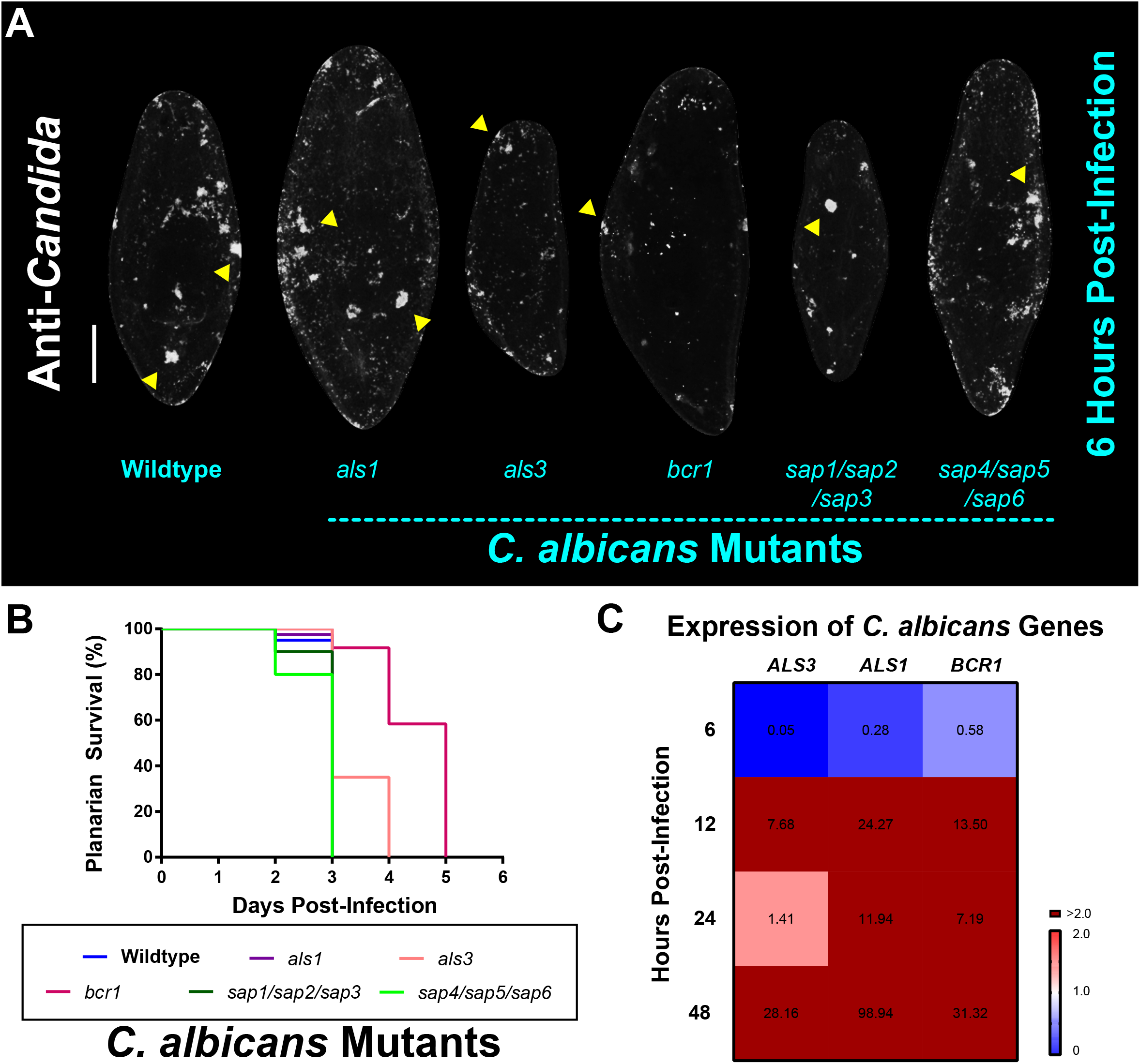
*C. albicans* Als3 and Bcr1 are important for virulence in planarians. (A) Whole-mount anti-*Candida* antibody stain of *C. albicans* mutant strains six hours post-infection. All mutant strains are capable of adhering to the planarian epithelial surface. (B) Planarian survival after infection with 25 million cells/mL of the different *C. albicans* mutant strains at the third day of infection. (C) Gene expression levels represented in a heat map of *C. albicans* genes *ALS3, ALS1*, and *BCR1* at the different infection timepoints. Gene expression is represented by fold change normalized to an uninfected control. Color scale depicts red as upregulation and blue as downregulation; burgundy demonstrates upregulation greater than twofold. Data were obtained in triplicate per experiment for at least two biological and technical replicates. Scale bar is 200μm.

### Als3 and Bcr1 are important players in the planarian mitotic response to C. albicans infection

Given the importance of *C. albicans* Als3 and Bcr1 for colonization of the planarian epithelial surface, we next wanted to know whether Als3 and Bcr1 play roles in the mitotic response of planarians. We found that six hpi, all *C. albicans* mutant and wildtype strains were capable of mounting a mitotic response in planarians; however, infection with the *als3* and *bcr1* mutant strains displayed significantly less mitotic proliferation in planarians than that of the wildtype strain (**Figure 3A-B**). These findings indicate that Als3 and Bcr1 are involved in mediating the planarian mitotic response.

**Figure 3:**
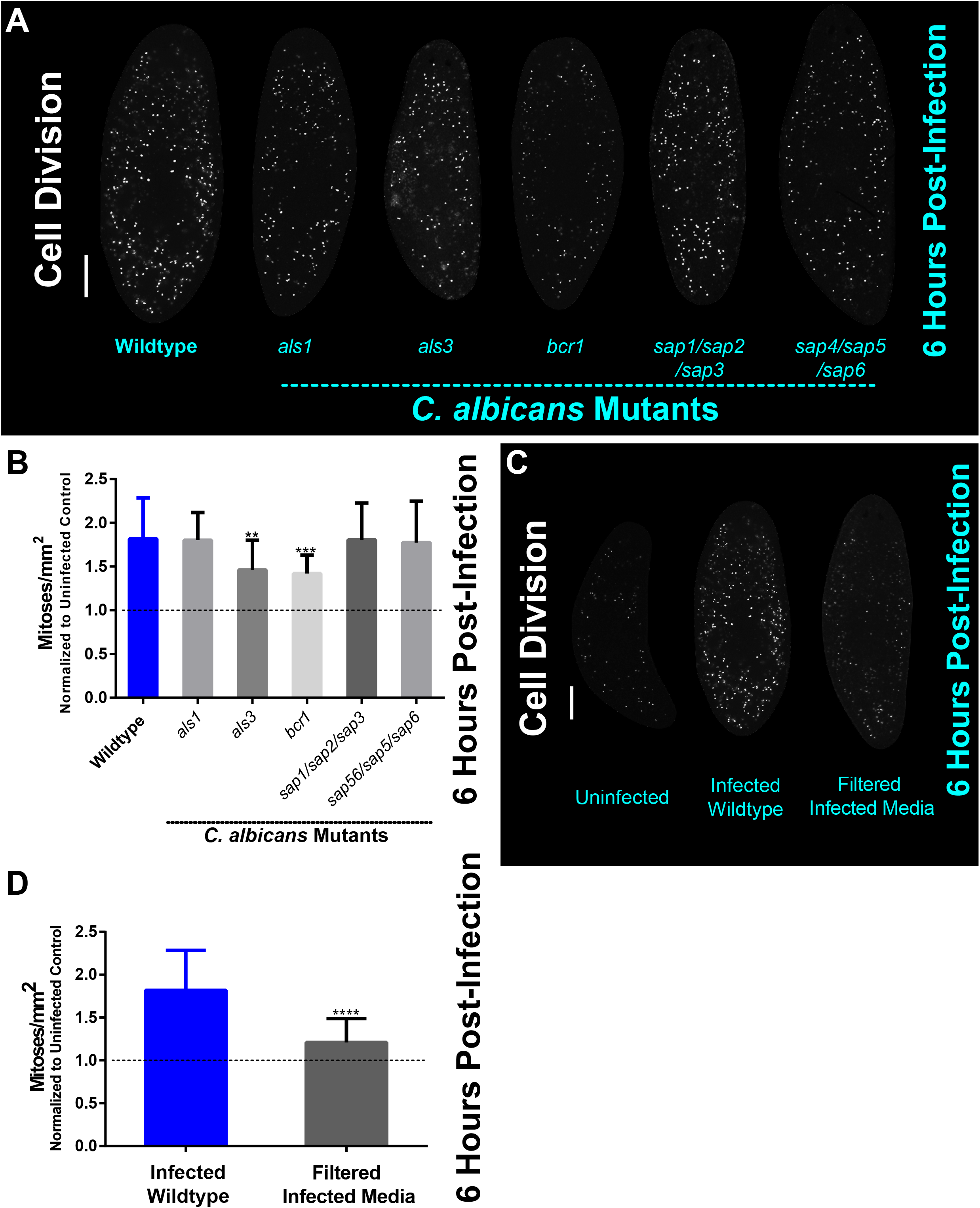
The planarian mitotic response decreases in infections with less virulent *C. albicans als3* and *bcr1* mutant strains. (A) Whole-mount immunostaining with anti-phospho-histone H3 (ser10) antibody for planarians, which labels mitotic cells (white foci) at six hours post-infection with the *C. albicans* mutant strains indicated. (B) Number of mitotic cells in response to *C. albicans* mutant strains normalized to uninfected planarians at six hours post-infection. C) Whole-mount immunostaining with anti-phospho-histone H3 (ser10) antibody for planarians, which labels mitotic cells (white foci) at six hours post-infection for animals that were soaked in filter sterilized cell-free infected planarian media. (D) Whole mount immunostaining with anti-phospho-histone H3 (ser10) antibody, which labels mitotic cells (white foci) at six hours post-infection normalized to an uninfected control. *C. albicans* infections were performed using 25 million cells/mL. Graphs represent mean ±SEM. All statistical comparisons are against the wildtype strain unless noted with bars. Scale bar is 200μm. Two-way ANOVA, **P<0.005; ***P<0.001; ****P<0.0001.

Since secreted aspartyl proteases (Saps) are important *C. albicans* virulence factors, we hypothesized that Saps may play roles in the planarian host response to *C. albicans* infection. To address this, we exposed planarians to cell-free filter sterilized media obtained from six hpi infection assays and measured planarian mitotic activity. We found no difference in the mitotic activity between uninfected animals and the group exposed to filter sterilized media from the infection assays, suggesting that the increase in cell proliferation upon *C. albicans* infection requires the physical presence of the pathogen (**Figure 3C-D**). Altogether, these findings suggest that adherence of *C. albicans* cells to the planarian epithelial surface, rather than fungal secreted proteases, induces the host mitotic response during the early hours post-infection.

### The planarian Dectin signaling pathway modulates the early mitotic response to and the clearance of the fungal infection

In order to understand how *C. albicans* cells interact with the planarian host to trigger hyper-proliferation, we decided to explore the initial molecular response mediated by the planarian innate immune system upon infection. Our previous work showed that the expression of genes in the host Dectin signaling pathway are generally upregulated one day post-infection with *C. albicans* (Maciel et al., 2019). To expand on these initial findings, we screened, by qRT-PCR, an expanded list of host genes encoding components of the Dectin signaling pathway, including the gene encoding the upstream SYK adapter protein *Smed-syk (syk)* and downstream gene components *Smed-card* (*card*), *Smed-bcl* (*bcl*), *Smed-malt1* (*malt1*), *Smed-tab1* (*tab1*), and *Smed-tak* (*tak*) during early infection with *C. albicans* (Wagener et al., 2018). We found that as early as three hpi there was a significant increase in the expression of all genes assayed of the Dectin signaling pathway, and that the upregulation of each gene was generally sustained for 48 hpi (**Figure 4A**). Notably, the upstream gene *syk* displayed an increase in expression (~threefold) in the first 6-12 hpi. We confirmed this upregulation in expression using fluorescent *in situ* hybridization (FISH) with a probe against *syk*, which revealed *syk* expression throughout the animal with enrichment in the digestive system including the pharynx of uninfected planarians (**Figure 4B**). In the infected planarians, we observed that *syk* expression became more prominent with the appearance of punctate foci readily visible in the main intestinal branches close to the brain at six hpi (observed in 6 out of 8 animals).

**Figure 4:**
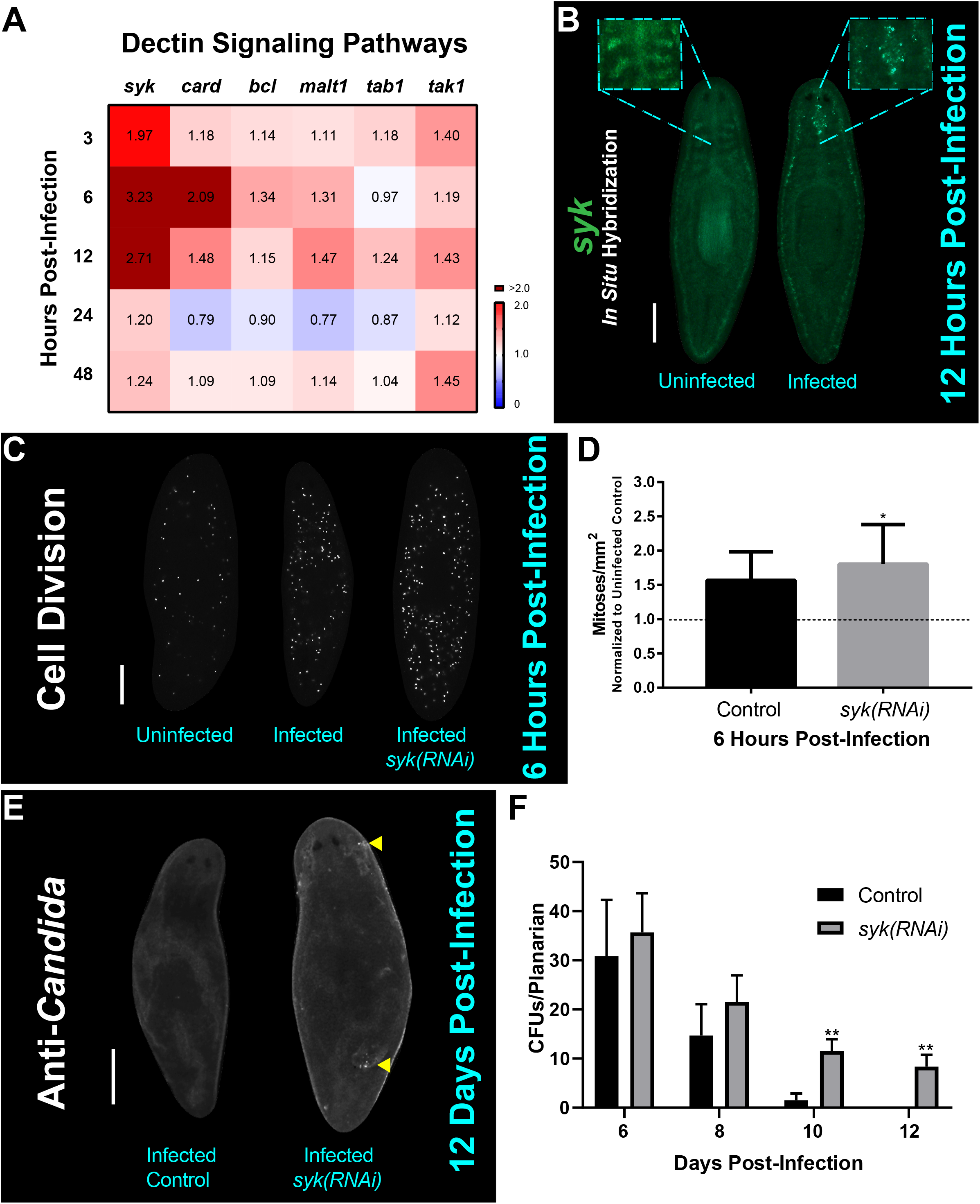
The Dectin signaling pathway plays a role in the host response to and clearance of *C. albicans* infection. (A) Gene expression levels for Dectin signaling pathway homologs at early and late timepoints of *C. albicans* infection using 25 million *C. albicans* cells/mL. Gene expression is represented in a heat map displaying fold change normalized to an uninfected control. Color scale depicts red for upregulation and blue for downregulation; burgundy demonstrates upregulation over twofold. Data were obtained in triplicate per experiment for at least two biological and technical replicates. (B) Fluorescent *in situ* hybridization showing expression of *syk* in an uninfected and twelve-hour infected animal. The boxed regions are magnifications of representative uninfected and infected animals. This experiment was replicated two times using five animals per experiment. Scale bar is 200μm. Images are representative of eight animals for at least two biological and technical replicates. (C) Whole mount immunostaining with anti-phospho-histone H3 (ser10) antibody, which labels mitotic cells (white foci) at six hours post-infection. (D) Number of mitoses of control infected animal versus *syk(RNAi)* infected animals normalized to an uninfected control at six hours post-infection. (E) Representative images of whole mount anti-*Candida* antibody stain at twelve days post-infection (white foci) of a control animal and *syk(RNAi)* animal. (F) Number of *C. albicans* colony forming units (CFUs) normalized per planarian of infected control versus infected *syk(RNAi)* planarians throughout the course of infection. This experiment was replicated two times using five animals per experiment. 25 million *C. albicans* cells/mL was used for the infections. All graphs represent mean ±SEM. Scale bar is 200μm. T-test, *P<0.01; **P<0.001.

To assess the role of the Dectin signaling pathway in the planarian innate immune response, we disrupted *syk* function with RNA-interference (RNAi) and evaluated the planarian mitotic response and ability to clear the *C. albicans* infection. Intriguingly, downregulation of *syk* by RNAi led to a slight increase (13%) in mitotic activity when compared to the infected control at six hpi (**Figure 4C-D**). Loss of *syk* function by RNAi resulted in planarians having a reduced capacity to clear the infection, even after 12 days post-infection (**Figure 4E-F**). Taken together, these findings suggest that the Dectin signaling pathway is an important mediator of the innate immune response in planarians that modulates the early mitotic response to fungal infection and is required for proper clearance of the infection.

### C. albicans infection triggers a heterogenous transcriptional response across planarian neoblast subpopulations

Recent work suggested a model in which transcriptomic changes occur in planarian neoblasts as they adopt new cellular identities necessary to support cellular turnover and tissue regeneration (Zeng et al., 2018). To gain a deeper understanding of how neoblast dynamics are affected in response to *C. albicans* infection, we screened, by qRT-PCR, planarian genes encoding neoblast markers for various subpopulations of progenitor cells. We measured changes in gene expression for six different neoblast lineages at different timepoints post-infection, and our results revealed that transcriptomic changes were not homogenous throughout the neoblast subpopulations. We discovered an upregulation in the expression of several neoblast subclasses detected as early as three hpi and this upregulation was maintained for the next 48 hpi (**Figure 5A**). The neoblast lineages measured represent twelve neoblast clusters of cells (NB1-NB12) and are ordered based on *Smed-piwi-1 (piwi-1)* transcript levels; thus, NB1 expresses higher levels of *piwi-1* relative to NB12 (Zeng et al., 2018). We found that in the first 24 hpi there was a reduction in the expression of *Smed-tspan-1 (tspan-1)*, which defines the NB2 subpopulation that contains the pluripotent clonogenic neoblasts (cNeoblasts) (**Figure 5A**) (Wagner et al., 2011; Zeng et al., 2018). The expression of *tspan-1* was slightly increased at 48 hpi. We also found that expression of the *Smed-lmo3 (lmo3)* marker (NB3) was consistently downregulated throughout the first 48 hpi. Overall, we detected the highest expression levels associated with markers of the neoblast subpopulations NB9, NB11, NB4, NB7, and NB5. The absolute highest expression levels were detected with the marker *Smed-pou2-3* (*pou2-3*) (NB9), which gradually increased after three hpi and peaked more than eightfold at 48 hpi (**Figure 5A**). High expression levels were also detected for markers *Smed-ston2* (*ston2*) and *Smed-pdch11 (pdch11)* (NB11 and NB4, respectively). Of note, *Smed-pj-1b (pj-1b)* of the NB4 neoblast subpopulation was the only marker whose expression increased dramatically while the other eight NB4 markers remained close to the levels of the control (**Figure 5B**). Altogether, these results suggest that infection with *C. albicans* triggers a heterogenous transcriptional response across planarian neoblast subpopulations.

**Figure 5:**
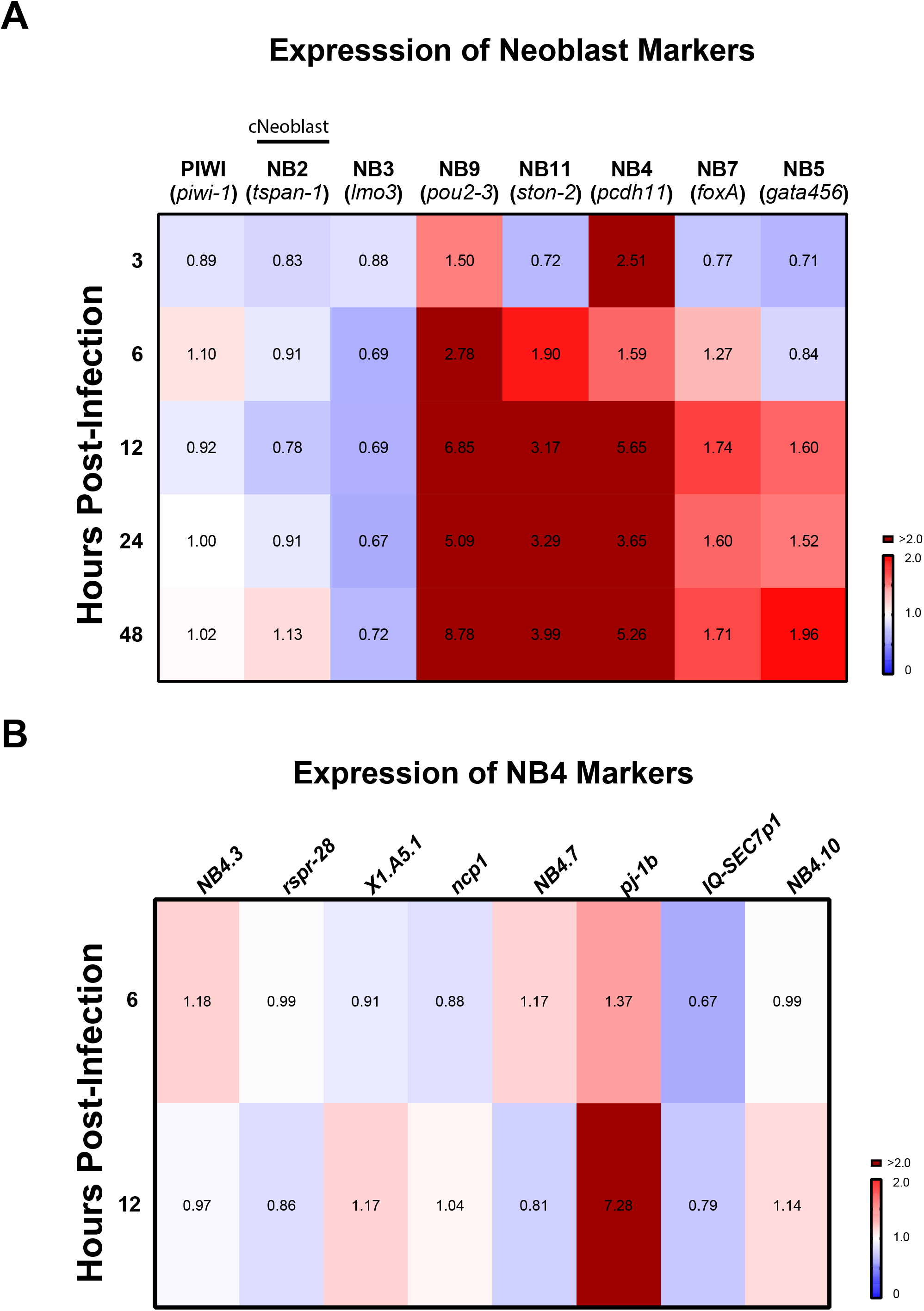
Expression of planarian neoblast subclasses throughout infection with *C. albicans*. (A) Gene expression levels of neoblast markers and subclass markers of clonal neoblasts over the course of *C. albicans* infection. Expression levels are represented in a heat map, where the color scale depicts red as upregulation and blue as downregulation; burgundy is upregulation greater that twofold. Gene expression is represented by fold change normalized to an uninfected control. Gene expression values represent triplicate samples for at least two biological and technical replicates. B) Gene expression of eight other markers for NB4 (Zeng et al., 2018).

### C. albicans infection modulates transcription within neuronal clusters and the excretory system of planarians

To further examine how increased gene expression levels of the NB9 and NB11 neoblast subpopulations impact the planarian response to infection with *C. albicans*, we extended our analysis to include markers of downstream planarian progenitor cells that lead to differentiated cells in the excretory and nervous systems (Rink et al., 2011; Ross et al., 2017; Zeng et al., 2018). The early progenitors of the nervous (NB11) and excretory (NB9) systems were upregulated during infection with *C. albicans* (Figure 5A), suggesting that the planarian infection response goes beyond neoblast subpopulations to also involve post-mitotic committed cells.

The planarian central nervous system is comprised of distinct neural subtypes, labeled by the synthesis of neurotransmitters that they produce (Ross et al., 2017). These subtypes include dopaminergic, octopaminergic, GABAergic, serotonergic, and cholinergic neurons, which all have specific markers (*Smed-th (th), Smed-tbh (tbh), Smed-gad (gad), Smed-tph (tph)*, and *Smed-chat (chat)*, respectively). We wanted to understand how the increase in expression of NB11 upon *C. albicans* infection would impact the different neuronal subpopulations. To explore this idea, we measured the expression levels, by qRT-PCR, of these six neural markers at both 12 and 24 hpi, which were the timepoints where the expression of NB9 and NB11 were highest (**Figure 5A**). We found that the expression levels for two neuron subtypes, *tbh* and *chat* were increased when compared to uninfected animals, with *chat* displaying the highest levels of upregulation (**Figure 6A**). We confirmed our qRT-PCR results by performing FISH using a *chat* probe, which is expressed along the brain and the ventral cords in uninfected planarians. The *chat* signal was found to be increased in the brain of infected planarians starting at three hpi (**Figure 6B**). We also found that the increase in expression of *chat* and the downstream adapter gene *syk* of the Dectin signaling pathway can be detected as early as 15-30 minutes post-infection and remains elevated for the first three hpi (**Figure 6C**). The expression levels of both *chat* and *syk* were observed to increase at 30 minutes and continued to increase in the later timepoints of infection.

**Figure 6:**
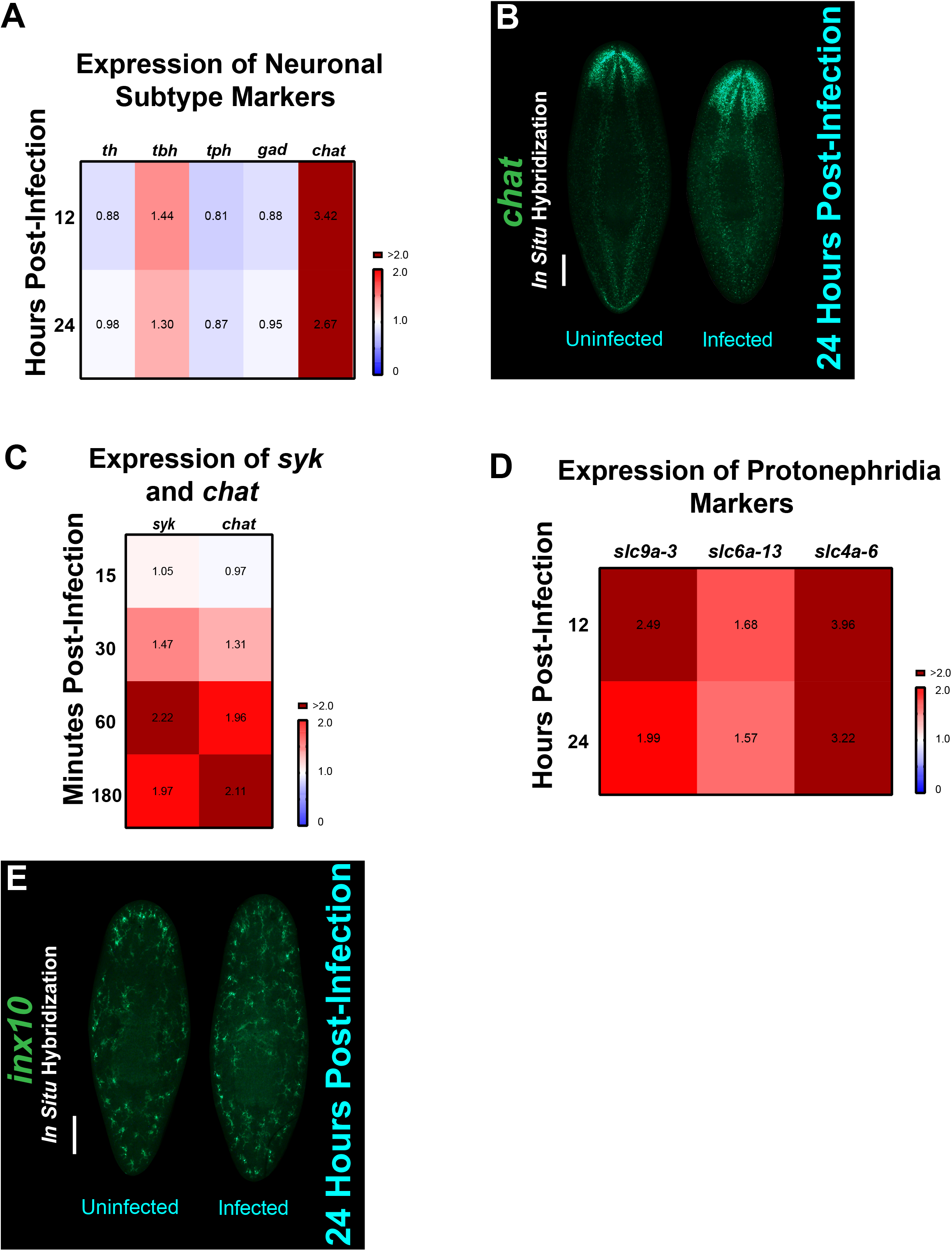
Neurons, protonephridia, and specific mesenchymal neoblast markers are upregulated in planarians during infection with *C. albicans*. A) Expression of five different neuron subtypes. Gene expression levels throughout the course of the *C. albicans* infection are represented in the heat map, where the color scale depicts red as upregulation and blue as downregulation; burgundy is upregulation greater than twofold. Gene expression is represented by fold change normalized to an uninfected control. B) Fluorescent *in situ* hybridization showing expression of *chat* in an uninfected and 24-hour infected planarian. Scale bar is 200μm. Images are representative of eight animals for at least two biological and technical replicates. C) Gene expression levels of *syk* and *chat* throughout the course of the *C. albicans* infection (15-180 minutes). Expression levels are represented in a heat map, where the color scale depicts red as upregulation and blue as downregulation; burgundy is upregulation greater than twofold. Gene expression is represented by fold change normalized to an uninfected control. Gene expression values represent triplicate samples for at least two biological and technical replicates. D) Expression of protonephridia structure markers: *slc9a-3* (collecting ducts), *slc4a-6* (distal tubules), and *slc6a-13* (proximal tubules). Gene expression levels throughout the course of the *C. albicans* infection are represented in the heat map, where the color scale depicts red as upregulation and blue as downregulation; burgundy is upregulation greater than twofold. Gene expression is represented by fold change normalized to an uninfected control. E) Fluorescent *in situ* hybridization showing expression of *inx10* in an uninfected and 24-hour infected animal. Scale bar is 200μm. Images are representative of eight animals for at least two biological and technical replicates.

To determine how the planarian excretory system is impacted by the increase in NB9 expression, we measured markers of the protonephridia, and more specifically, markers of the flame cells that contribute to electrolyte balance and mucus production, among other excretory functions (Rink et al., 2011). We observed an increase in expression of the markers of collecting ducts *Smed-slc9a-3 (slc9a-3)*, distal tubules *Smed-slc4a-6 (slc4a-6)*, and proximal tubules *Smed-slc6a-13 (slc6a-13)* during the first 24 hpi (**Figure 6D**). We validated these results by performing FISH using a *Smed-inx10 (inx10)* probe (Oviedo and Levin, 2007), which confirmed a steady increase in *inx10* expression throughout the planarian body at 24 hpi (**Figure 6E**). Overall, these findings suggest that both the nervous and excretory systems contribute to the host response during fungal infection.

## DISCUSSION

Here, we show that the planarian infection model is an advantageous animal model system to study host-pathogen interactions, particularly at early stages of epithelial infection with *C. albicans*. Our results demonstrate that the superficial adherence of *C. albicans* to the planarian epithelial surface triggers a multi-system host response involving stem cells and differentiated tissues in the planarian **(Figure 7)**.

**Figure 7:**
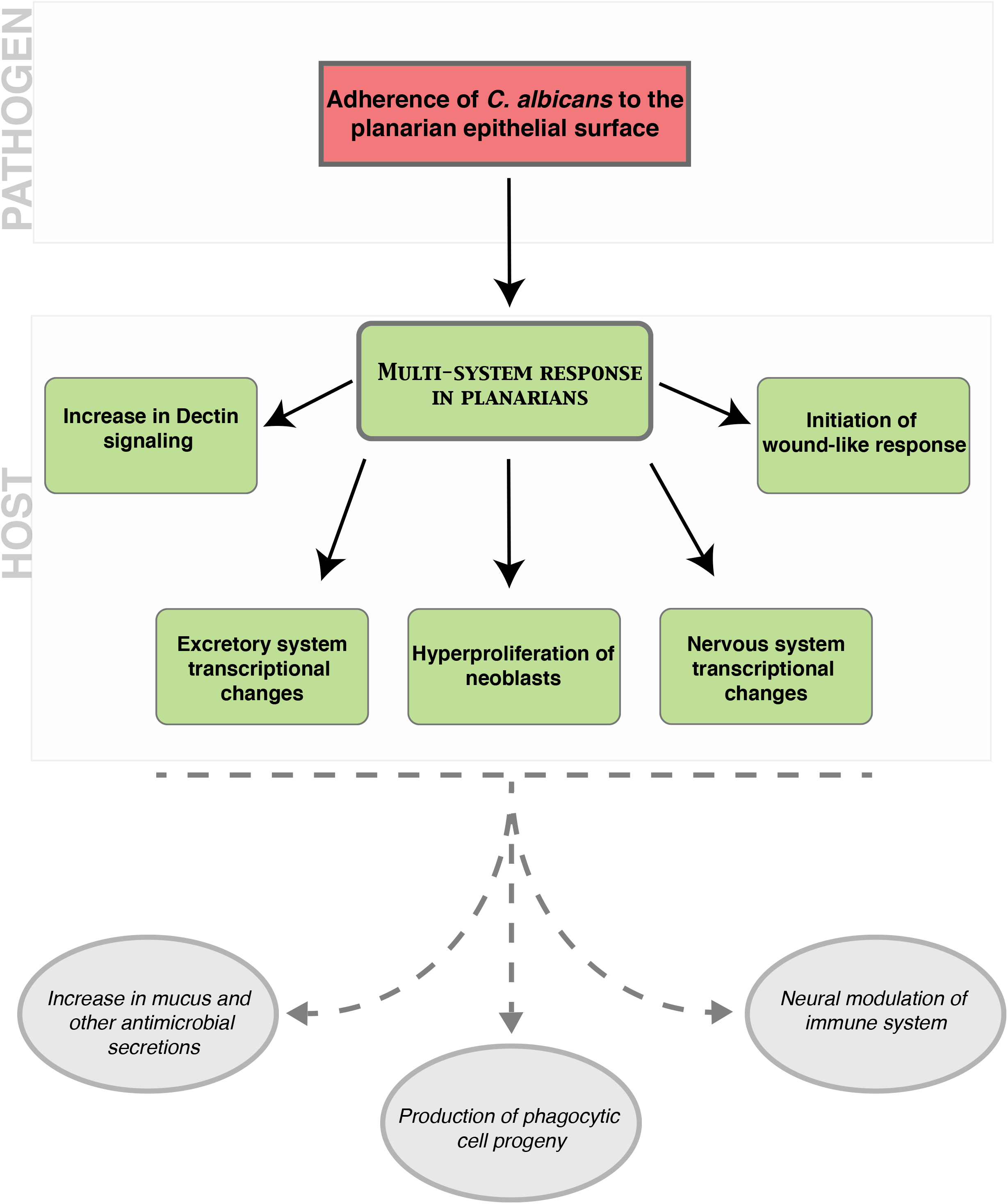
Model of the cascade of host-pathogen interactions that occur following an epithelial infection of planarians with *C. albicans*. The top red box indicates the role of *C. albicans* in initiating a multi-system response in planarians during an infection. The middle green boxes indicate the planarian responses to *C. albicans* during an infection including an increase in components of the Dectin signaling pathway, the initiation of a wound-like response, the hyperproliferation of neoblasts, and transcriptional changes in the excretory and nervous systems. The bottom grey ovals represent inferred host outcomes of the planarian multi-system response to the *C. albicans* infection. We propose that an increase in mucus and other antimicrobial secretions, the production of phagocytic cell progeny, and neural modulation of the immune system are downstream outcomes of this host-pathogen interaction.

These findings underscore the advantages of analyzing host-pathogen interactions *in situ* as a powerful paradigm to identify crosstalk between tissues during fungal infection. Planarians are thus a unique model system to study the host response to infection at the organismal level, a feature that is largely missing from or challenging to achieve with existing animal models of infection.

Our results reveal that part of the host response to *C. albicans* epithelial infection involves a stereotypical pattern of neoblast division. Two systemic peaks of neoblast division occur at six and 24 hpi without visible signs of host tissue damage. This is intriguing because physiological increases in neoblast division generally occur via metabolic inputs such as nutrient availability through feeding or in response to tissue injury (Peiris et al., 2012; Saló and Baguñà, 1984; Wenemoser and Reddien, 2010).

The metabolic-induced neoblast hyperproliferation is systemic and tends to peak around 6-12 hours post-feeding to gradually return to pre-feeding levels by 48 hours (Baguñà, 1976; Kang and Sánchez Alvarado, 2009). The amputation-induced neoblast division takes place as biphasic peaks that occur first systemically at six hours post-amputation and then localized near the injury site between 48-72 hours following wounding (Wenemoser and Reddien, 2010). Our findings suggest that similar to amputation, fungal infection triggers neoblast proliferation by six hpi and it is preceded by ERK phosphorylation and the overexpression of the early wound response gene *egr2* (Owlarn et al., 2017; Wenemoser and Reddien, 2010). However, the neoblast response to *C. albicans* infection does not involve an increase in the expression of *runt1*, which is important for regeneration, and is not triggered by major tissue damage as observed by amputation or the resulting distention of the intestine that occurs after feeding.

Furthermore, the second mitotic peak observed after fungal infection is systemic and occurs roughly 18 hours after the initial peak. Another striking difference with the regeneration response is that cell death does not precede the neoblast increase in proliferation during the early stages of infection (**Supplemental Figure 2**). Therefore, the mitotic response to fungal infection differs in time and location with respect to other stimuli such as feeding and amputation. We speculate that *C. albicans* infection activates an infection-specific neoblast proliferative response.

The neoblast hyper-proliferative response is preceded by the adherence of *C. albicans* to the planarian epithelial surface. This response is dependent on the *C. albicans* transcription factor Bcr1, which is a major regulator of adherence, and its downstream GPI-anchored protein Als3, which is an important adhesin. Although the precise molecular mechanisms leading to hyper-proliferation in planarians in response to infection with *C. albicans* are unclear, it seems likely that the ability of *C. albicans* to adhere to the planarian epithelial surface is a critical factor in inducing this response. Given that Bcr1 controls the expression of numerous genes encoding cell wall proteins, and is a major regulator of biofilm formation (Nobile et al., 2006; Nobile et al., 2012; Nobile and Mitchell, 2005), it seems likely that the ability to colonize and form a biofilm on the planarian epithelial surface are important fungal processes in initiating this hyper-proliferative host response. Als3 is also required for biofilm formation in *C. albicans* and is a major adhesin expressed on the surface of hyphal cells. Thus, the ability to form adhesive hyphae that can actively penetrate host epithelial cells is another factor that likely mediates this host response (Wachtler et al., 2011). Interestingly, Als3 is known to bind to mammalian host ligands, such as E-cadherin on epithelial cells and N-cadherin on endothelial cells (Phan et al., 2005; Phan et al., 2007), which induces engulfment of fungal cells into the mammalian host via a clathrin-dependent mechanism. Given the importance of Als3 in triggering hyper-proliferation in planarians, it is possible that the process of host cell engulfment of *C. albicans* cells is also involved in initiating this host response. Taken together, it seems likely that both invasion mechanisms of induced endocytosis as well as active penetration of *C. albicans* cells are complementary mechanisms that induce hyper-proliferation of the planarian host.

Adherence of *C. albicans* cells to the planarian epithelial surface is followed by a generalized overexpression of host genes associated with the Dectin signaling pathway. In our previous work (Maciel et al., 2019), we also observed an overexpression of components of the Dectin signaling pathway (*syk* and *tak*) at later timepoints of the infection, suggesting that there is a consistent innate immune response during fungal infection. A key component of the Dectin signaling pathway is the adaptor protein SYK, and our results show a persistent overexpression of the *syk* gene throughout infection with *C. albicans*. Unexpectedly, functional disruption of *syk* by RNAi resulted in a slight increase in neoblast proliferation upon *C. albicans* infection. It remains unclear why this apparent increase in neoblast proliferation is accompanied by a delayed clearance of the infection. It is possible that this is due to redundancy with other pathways or to an aberrant neoblast response to infection. Nonetheless, these findings suggest that *syk* is required for planarians to clear the *C. albicans* infection in a timely manner.

As part of the host response to *C. albicans* infection, we observed a differential expression in planarian markers of neoblast subpopulations. This finding suggests that not all neoblasts are engaged in the response to fungal infection and opens up the possibility of an infection-specific neoblast response. Recent work classified neoblast subpopulations based on the content of the marker *piwi-1* (Zeng et al., 2018). This study found that neoblasts with high *piwi-1* content (*piwi-1*^*high*^) include the clonogenic neoblasts (cNeoblasts) that together are key players in the early regenerative response (Zeng et al., 2018). However, we found that the expression of markers for neoblast subpopulations with the highest content of *piwi-1* (e.g., NB1, NB2, and NB3) remained relatively low or did not change relative to the uninfected group in the initial 48 hpi.

Instead, there was a dramatic increase in the expression of markers for neoblasts with reduced levels of *piwi-1* (i.e., NB4, NB5, NB7, NB9, and NB11) upon *C. albicans* infection. From these neoblasts subclasses NB9, NB11, and NB4 displayed the highest expression levels, greater than eightfold in the initial 48 hpi. These three neoblast subpopulations contain less than 5% of *piwi-1* positive cells (Zeng et al., 2018). This finding raises the possibility that the active cycling of neoblasts with low *piwi-1* content could be responsible for the two mitotic peaks we observed during infection with *C. albicans*. We propose that *C. albicans* infection elicits a heterogenous neoblast response that is distinct from that observed during regeneration.

Our findings indicate that the host response to *C. albicans* infection involves transcriptional changes in distinct differentiated planarian tissues. The upregulation in the expression of genes associated with the excretory system is detected as early as 12 hpi. The increased expression of genes within the excretory system may lead to an enhanced secretion of mucus, which contains antimicrobial components to aid in the elimination of the pathogen. The mucus barrier in planarians contains an extensive array of antimicrobial peptides, zymogens, and proteases that aid in the innate immune response to infection (Bocchinfuso et al., 2012). Likewise, the dramatic increase in the expression of the marker of cholinergic neurons (*chat*) in the initial 24 hpi, suggests that this neuronal type may be important in the innate immune response to *C. albicans* infection. This is consistent with the idea that certain neuronal groups can act as immunocompetent cells, which is an ancient function conserved between pre-bilaterians and mammals (Godinho-Silva et al., 2019; Klimovich et al., 2020). Indeed, murine nociceptive neurons are critical for the innate immune response to cutaneous infection with *C. albicans* (Kashem et al., 2015). Together, the increase in expression of markers of the neural and the excretory systems in planarians suggests the existence of long-range organismal communication that is likely needed to orchestrate defense against invading pathogens. It is tempting to speculate that the adherence of *C. albicans* cells to the planarian epithelial surface may trigger neural cues that activate the secretion of mucus with antimicrobial properties and the innate immune response through the Dectin signaling pathway. Future experiments will address the precise interplay between neoblasts, differentiated tissues and the molecular cascade orchestrated by the planarian innate immune system to defend against invading pathogens **(Figure 7)**.

## MATERIALS AND METHODS

### Planarian culture

The planarian colony used for all assays was CIW4, an asexual *Schmidtea mediterranea* strain. The colony was maintained as previously described (Oviedo et al., 2008b).

### Microorganisms

The *Candida albicans* wildtype strain SN250, a derivative of the clinical isolate SC5314, was used as the isogenic wildtype strain along with the *bcr1* mutant strain (TF137) (Homann et al., 2009); the *als1* mutant strain and the *als3* mutant strain (Nobile et al., 2008); and the triple *sap1/sap2/sap3* mutant strain and the triple *sap4/sap5/sap6* mutant strain (Felk et al., 2002). All *C. albicans* strains were grown from cryogenic stocks on yeast extract peptone dextrose (YPD) agar for 48 hours at 30°C. Single colonies were then inoculated into liquid YPD media and grown for 12-6 hours at 30°C for use in the infection assays.

### Infection assays

Ten animals were kept in 6mL wells with 4mL of planarian water (media) containing 25 million *C. albicans* cells per milliliter as previously described (Maciel et al., 2019). The animals were kept in the infected media until collected for specific experiments at various timepoints. The infection assay took place at 25°C, the preferred temperature for planarians, for the duration of the experiments.

### *C. albicans* CFU measurements

Planarians were collected at the indicated timepoints during the infection and rinsed with planarian media. The animals were homogenized in 500μL of planarian media and diluted in 10mL of planarian media. After homogenization, 300μL of the homogenate was plated onto YPD media agar plates containing a cocktail of broad-spectrum antibiotics. The colonies were counted to obtain CFUs after being incubated at 30°C for 48 hours.

### Protein extractions

Animals were washed thoroughly prior to being placed in 1.5mL centrifuge tubes. Approximately 150μL of 1X RIPA Buffer (Cell Signaling Technologies, 9806) containing protease inhibitors (Complete Mini Protease Inhibitor Cocktail (Roche, 04693124001); Halt Phosphatase Inhibitor cocktail (Thermo Scientific, 1862495), 1mM PMSF, 1mM DTT) was added to each tube. Samples were placed on ice and homogenized using a motorized pestle (Fisher,12-141-361) for 10 seconds, followed by a 45-minute incubation. Samples were then centrifuged at 20,817 g for 20 minutes at 4°C. Approximately 100μL of supernatant was transferred into a clean tube and placed on ice. Protein concentrations were determined using a Bradford protein assay (VWR, E530-1L). In a 1.5mL tube, 50μg of supernatant was mixed with 6X Laemmli buffer (6%SDS,9% β-mercaptoethanol,4.8% glycerol, 0.03% bromophenol blue, 375mM Tris-HCl) and was heated to 94°C for 10 minutes to denature and reduce the proteins.

### Western blots

Protein lysate mixtures were loaded onto a 15% SDS-PAGE gel along with a molecular weight marker (Thermo Scientific, 26619). Samples were transferred to a methanol-activated PVDF membrane (Bio-Rad,162-0175) for 2 hours at 55 V in a 1X Tris-glycine transfer buffer (25mM Tris base, 192mM glycine,10%(v/v) methanol) on ice. The membrane was blocked with 5% BSA in TBST (20mM Tris-base, pH 7.6, 140mM NaCl, 0.1%Tween-20) for 2 hours at room temperature and incubated with the primary antibody dilutions for 16 hours at 4°C on a rocker. The primary antibodies used were anti-actin (1:3000; Developmental Studies Hybridoma Bank, JLA20), anti-pERK (1:400; a gift from the Bartscherer lab) (Owlarn et al., 2017). The primary antibody was removed and the membrane was washed with TBST four times (5 minutes each wash), before the addition of the secondary antibody: goat-anti-rabbit HRP IgG antibody (1:10,000; Abcam, ab7097) for anti-ERK, goat-anti-mouse HRP IgG antibody (1:10000; Invitrogen, G-21040) for anti-actin. Secondary antibody dilutions were incubated for 1 hour at room temperature with 5% non-fat milk in TBST-SDS (20mM Tris-base, pH 7.6, 140mM NaCl, 0.2%Tween-20, 0.01% SDS). The membrane was washed with TBST four times (5 minutes each wash), followed by two 5-minute washes with 1X PBS. The addition of the HRP chemiluminescence substrate (Millipore, WBLUF0100A) allowed for the detection of a signal. For the removal of primary antibody pERK, blots were stripped for 15 minutes (Thermo Scientific, 21059) and washed with 1XPBS four times (5 minutes each wash) before blocking for actin. Band intensities were quantified by computing the area under the curve using imageJ software bundled with Java (Schneider et al., 2012). pERK activity was then normalized to actin. Blots were imaged using a ChemiDoc gel imaging system (Bio-Rad).

### Whole-mount immunofluorescence

Non-infected and infected planarians were sacrificed with 5.7% 12N HCL solution for 5 min and fixed using Carnoys solution for 2h on ice. After the fixation, animals were stored in methanol at −20°C and then bleached overnight in a 6% H_2_O_2_ solution. Animals were then rehydrated in dilutions of methanol:PBSTx and stained as previously described (Ziman et al., 2020). The primary antibody used was α-H3p, 1:250 (Millipore Cat# 05–817R). The secondary antibodies used were goat anti-rabbit Alexa568, 1:800 (Invitrogen Cat# 11036) and HRP-conjugated goat anti-rabbit antibody (Millipore Cat# 12-348).

To stain *C. albicans* cells, infected animals were sacrificed in 10% NAC diluted in PBS. The planarians were then fixed in 4% formaldehyde in PBTx and permeabilized in 1% SDS. The animals were bleached in 3% H_2_O_2_ in 1X PBS. The primary antibody used was anti-*Candida*, 1:500 (ThermoFisher Cat# PA1-27158). The secondary antibody used was goat anti-rabbit Alexa568, 1:800 (Invitrogen Cat# 11036).

### RNAi treatments

Double-stranded RNA (dsRNA) was synthesized as previously described (Oviedo et al., 2008a). The dsRNA was applied via microinjections in three consecutive days with a fourth injection a week after the last injection. The animals were starved for at least one week prior to the dsRNA injections.

### Whole-mount fluorescence *in situ* hybridizations

Riboprobes were made using T3 and T7 polymerases and a digoxigenin-labeled ribonucleotide mix (Roche, Cat.# 11277073910) using PCR templates, as described previously (Pearson et al., 2009). *In situ* hybridization with and without fluorescence was performed as described previously (King and Newmark, 2013).

### Imaging and data processing

Area measurements and cell counts were calculated using imageJ software bundled with Java (Schneider et al., 2012). The Nikon AZ100 Multizoom microscope was used to collect the digital images and the images were processed using NIS Elements AR3.2 software (Nikon). Photoshop software (Adobe) was used to adjust contrast and brightness.

### Gene expression analyses

RNA was extracted using TRIzol (Thermo Fisher Scientific). qRT-PCR reactions were performed using SYBR Green Master Mix in a 7500 Fast Real Time PCR cycler (Applied Biosystems). The TATA-box-binding protein domain gene was used as the internal control. Each experiment was performed in triplicate for each timepoint. qRT-PCR was performed as previously described (Peiris et al., 2012).

### Cell-free filter sterilized planarian media

The media from six hpi infection assays was removed from the planarian wells and filter sterilized using a Corning vacuum system with 0.22μm pore-size 13.6cm² PES Membrane (Cat. # 431153). Uninfected animals were then inoculated with the cell-free filter sterilized media using the infection assay described above.

### Statistical analyses

Two-way ANOVA or t-test statistics were performed, and data are shown as the mean ± SEM or fold change ± SEM unless otherwise noted. All statistics were performed by pooling biological replicates from each technical replicate using Prism7, Graphpad software Inc. (http://www.graphpad.com).

## Acknowledgements

We thank Edelweiss Pfister for lab management and planarian maintenance in the Oviedo lab, and Gurbinder Singh for technical assistance in the Nobile lab. We thank all members of the Nobile and Oviedo labs for insightful discussions and comments on the manuscript. We thank Dr. K. Bartscherer for kindly providing the pERK antibody. We thank Dr. Bernhard Hube for the gift of the *C. albicans sap1/sap2/sap3* and *sap4/sap5/sap6* triple mutant strains. The anti-actin antibody was obtained from the Developmental Studies Hybridoma Bank, created by the National Institutes of Health (NIH) National Institute of Child Health and Human Development (NICHD) and maintained at the University of Iowa, Department of Biology.

## Authors’ contributions

E.I.M., A.V.A, B.Z., C.J.N., and N.J.O. conceived, designed, and interpreted experiments. E.I.M., A.V.A, and B.Z. performed all experiments, acquired and analyzed data. E.I.M., A.V.A, B.Z., C.J.N., and N.J.O wrote the manuscript. All authors read the manuscript, provided comments and approved the final version.

## SUPPLEMENTAL FIGURE LEGENDS

**Supplement Figure 1:**
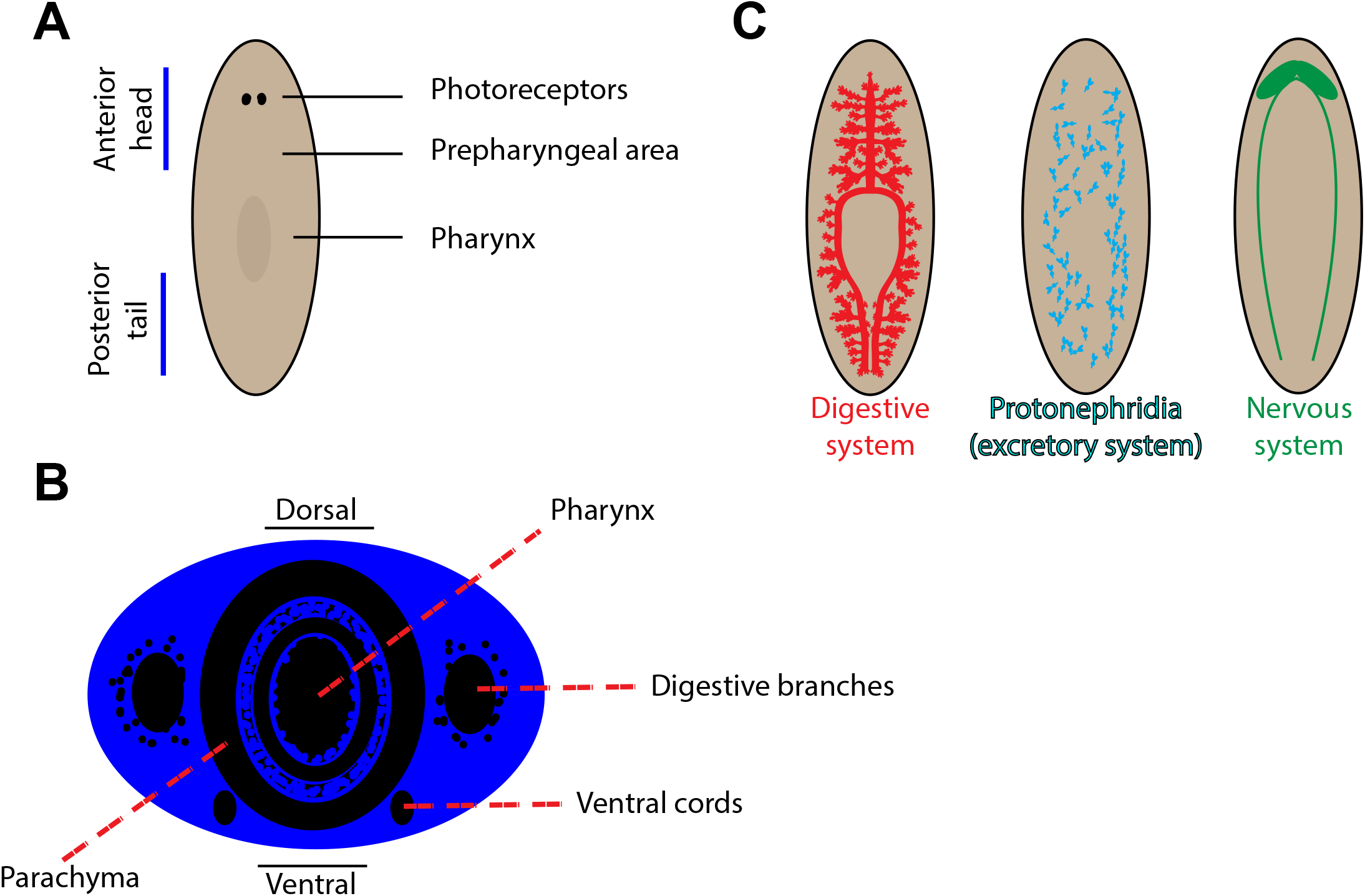
Orientation and basic anatomy of the planarian body. (A) Demonstrates the orientation the planarians in the whole-mount images. (B) Depiction of a transverse cross section of a planarian in the orientation found in the images throughout the figures. (C) Depiction of the digestive system, nervous system, and protonephridia of planarians.

**Supplement Figure 2:**
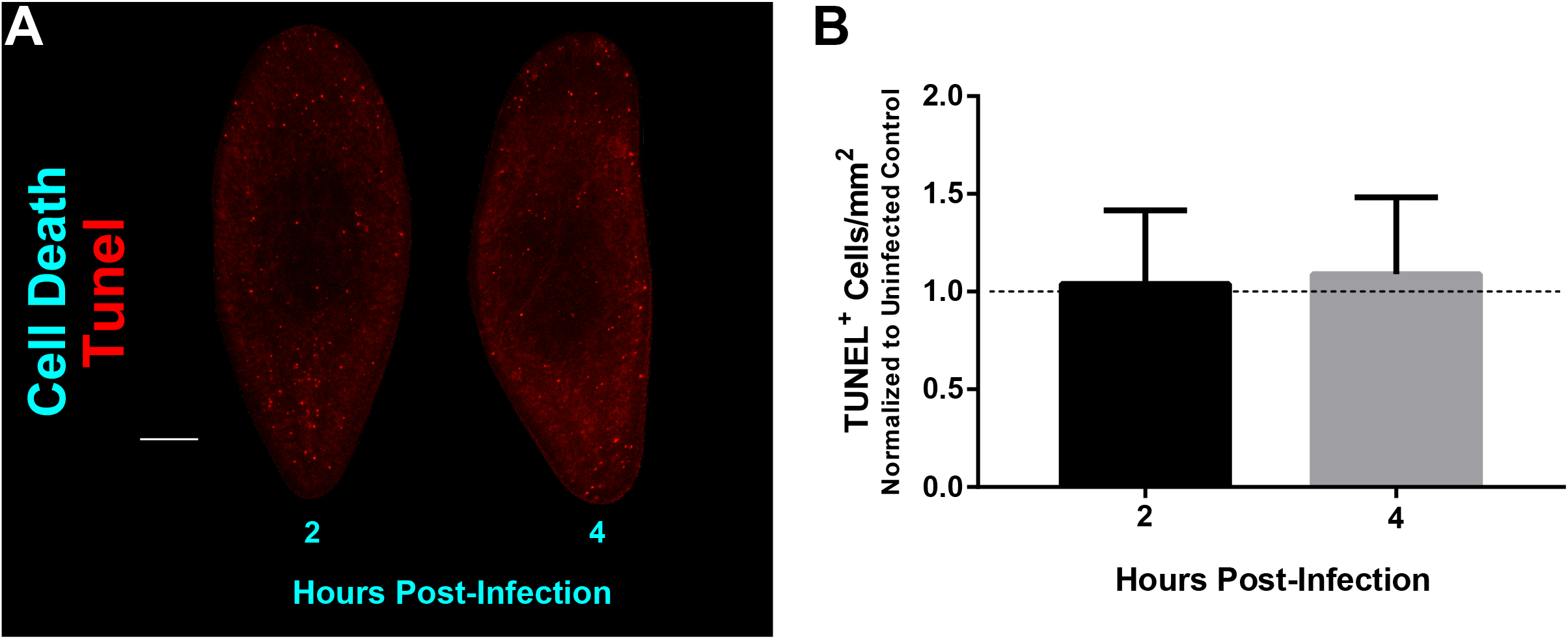
Cell death occurring during the early stages of infection with *C. albicans*. (A) TUNEL straining (red foci) was performed in uninfected planarians to compare animals infected at two and four hours post-infection using 25 million cells/mL of *C. albicans*. (B) Levels of TUNEL+ cells in the planarian tissue two and four hours post-infection normalized to an uninfected control. Cell death experiments consisted of two biological replicates using four animals each. All graphs represent mean ±SEM. Scale bar is 200μm.

